# Population receptive field tuning properties of visual cortex during childhood

**DOI:** 10.1101/213108

**Authors:** T. M. Dekker, D.S. Schwarzkopf, B. de Haas, M. Nardini, M.I. Sereno

## Abstract

Improvements in visuospatial perception such as contrast sensitivity and Vernier acuity continue until late in childhood, but the neural mechanisms driving these age-related changes are currently unclear. One contributing factor could be the protracted development of spatial tuning of neuronal populations across the visual cortex. Here we tested this possibility using population receptive field (pRF) mapping (Dumoulin and Wandell, 2008) in 6-to 12-year-old children and adults. We fitted pRF models to BOLD signals measured in areas V1-V4 and V3a during fMRI whilst participants watched wedge and ring stimuli traversing the visual field. Cortical magnification and the width of pRF tuning functions changed with viewing eccentricity in all participants. However, there were no age-related changes in pRF size, shape, cortical magnification, or map consistency across any of the visual areas measured. These results suggest that visuospatial perception in late childhood beyond age 6 years is not substantially limited by low-level spatial tuning properties of neuronal populations in visual cortex. Instead, performance improvements in this period may reflect more efficient use of the spatial information available in the visual system when forming perceptual judgments. These findings are an important step towards disentangling which neural mechanisms contribute to the eventual emergence of mature spatial vision, and for understanding the processes that determine the scope for visual plasticity at different stages of life.

## Introduction

Building a visual system with adult-like capabilities involves extensive shaping of neural mechanisms through experience (Hubel and Wiesel, 1962; Braddick and Atkinson, 2011). The most drastic changes in vision occur in the first year of life, but many visuospatial skills improve throughout the first decade. For example, contrast sensitivity matures between the ages of 7-9 years for gratings (Ellemberg et al., 1999; Adams and Courage, 2002; Benedek et al., 2003; Leat et al., 2009), and by age 10 for naturalistic textures (Ellemberg et al., 2009, 2012), whilst Vernier acuity (positional resolution) only converges on adult performance between10-14 years (Carkeet et al., 1997; Skoczenski and Norcia, 2002). Larger-scale spatial integration, such as acuity for shapes surrounded by flankers (crowding), size illusion sensitivity, and contour integration, continue to develop into the teenage years (Kovács, 2000; Doherty et al., 2010; Hadad et al., 2010; Jeon et al., 2010; Hipp et al., 2014).

What drives these substantial late changes in perception is an important unanswered question, especially since recent advances in eye disease treatments are increasing the need for understanding how age affects the plasticity of human visual pathways (*e.g.*, Carvalho et al., 2011). We used population receptive field (pRF) mapping (Dumoulin and Wandell, 2008) to test whether late childhood improvements in spatial vision reflect prolonged spatial tuning of neuronal populations across the visual hierarchy.

In the mature visual cortex, receptive fields of adjacent neurons are tuned to adjacent locations in the visual field, creating multiple retinotopic visual field maps (V1, V2, etc.). Receptive fields are smallest in neurons encoding the center of gaze, and more neurons are tuned to this part of the visual field. This adaptation is useful given the disproportionate cortical inputs from the fovea, where receptor density is highest and visual acuity greatest. In the living human brain, retinotopic tuning can be estimated at the voxel level using fMRI. A pRF model is fit to the BOLD response evoked by stimuli traversing the visual field (Dumoulin and Wandell, 2008). pRF tuning curves from human adults mirror those of single cell receptive fields; towards the periphery in visual cortex, pRF sizes increase (tuning functions become wider) and cortical magnification decreases (less cortex encodes the same visual distance). Recent studies have shown associations between individual adults’ pRFs and their Vernier acuity (Duncan and Boynton, 2003; Song et al., 2015), and found altered pRF tuning profiles in several populations with altered visuospatial processing (Brewer and Barton, 2014; Schwarzkopf et al., 2014; Clavagnier et al., 2015; Anderson et al., 2017). Thus, population-tuning curves in early visual cortex appear highly relevant for visuospatial skills.

Surprisingly little is known about visuospatial resolution in the developing human brain. Histological studies suggest that the fovea is still not mature by 45 months of age (~4 years) (Yuodelis and Hendrickson, 1986), and occipital cortex continues to thin and expand until ~age 10 years (Ducharme et al., 2016; Jernigan et al., 2016). However, the changes in neural functioning that accompany these structural changes are currently unclear. Here we investigate this question, by measuring pRFs across visual cortex in 6 to 12-year-old children and adults. If spatial tuning in visual cortex contributes to improved perception (*e.g.*, in Vernier- and crowded acuity) across these ages we may expect changes in *(i) pRF size*, as smaller pRFs are associated with greater visual acuity, and in *(ii) pRF shape components* that capture local interactions across neurons associated with crowding and context processing. We also expected to see changes in pRF distribution across the cortical sheet; specifically, in *(iii) cortical magnification*, since larger magnification is linked to greater visual acuity, and in *(iv) the consistency of retinotopic layout*, reflecting prolonged refinement of spatial maps that could facilitate the precision of position- and size information (Haak et al., 2013; Moutsiana et al., 2013). Alternatively, late improvements in visuospatial performance may reflect more efficient information read out by higher-level mechanisms, independent of spatial tuning in early visual cortex.

## Methods

### Subjects

We tested 39 children aged 6-12 years and 7 adults: all with normal or corrected to normal vision, no known neurological abnormalities. To ensure that any age-differences would not be driven by movement-related noise, we used stringent exclusion criteria; participants who made large movements (a 1 mm translation, or 3° rotation) during more than 3 volumes collected across all functional scans were excluded (see Data Quality Assurance and Supplementary Figure 1). This cut-off resulted in matched movement parameters across all age groups. The remaining participants included in the analysis were thirteen 6 to 9-year-old children (8.53 years, SD=0.89), seventeen 10- to 12-year-old children (Mean Age: 11.39, SD=0.74), and seven adults (22.30 years, SD=2.72). All participants had normal or corrected to normal vision, no known neurological abnormalities, met MR-safety criteria, and provided written informed (and in parental) consent. Experimental procedures were approved by the UCL Research Ethics Committee

### Scanning parameters

Structural and functional measures were obtained with a Siemens Avanto 1.5T MRI scanner and 30-channel coil (a customized 32 channel coil without obstructed view). BOLD measures were acquired using four single-shot EPI runs (TR=2.5s, volumes=144, slices=30 voxel size=3.2 mm^3^, axial plane, ascending, bandwidth=1930 Hz/pix, TE= 39 ms, flip=90). The high-resolution structural scan was acquired using a T1-weighted 3D MPRAGE (1 mm^3^ voxel size, Bandwidth=190 Hz/pix, 176 partitions, partition TR=2730, TR=8.4ms, TE=3.57, effective TI=1000 ms, flip angle=7 degrees).

### Stimuli

pRF mapping stimuli were back-projected on an in-bore screen at 34 cm viewing distance (projected area: 18 × 24 cm; resolution: 1920×1080). They consisted of moving ring and wedge checkerboards presented against a dark grey background. During odd runs, the ring expanded for 10 cycles (32 sec/cycle), and the wedge (21° angle) rotated anticlockwise for 6 cycles (53.33 secs/cycle). During even runs, the ring contracted and the wedge rotated clockwise at the same speeds. The stimuli covered a maximum vertical eccentricity of 14.8°, and moved to a new position each new TR. They were overlaid with a white central fixation dot (0.3 degree radius) and a white radial grid to anchor fixation. Checkerboards had a fixed contrast (35%), achieved by randomly selecting hue with constant saturation and two levels of lightness. Contrast reversals occurred at 8 Hz. Checkerboards were superimposed with moving dots (diameter: 0.08°; randomly rotating and expanding or contracting at speeds between 0°-19°/sec) and briefly presented photos of animals and household items (0.6 secs/image). Movement, color, and objects were added to elicit maximal retinotopic responses across visually driven cortex, and make the stimuli more appealing to children. The ring and wedge were preceded and followed by a 20-second (8 TR) baseline.

### Procedure

Participants “practiced” being scanned and lying still whilst watching a funny 5-minute cartoon and undergoing a short localizer scan. Then, participants completed 4 functional runs of 6 minutes with two “cartoon breaks” in-between when structural data were collected. To keep participants engaged and motivated to fixate during pRF mapping, they could score points by detecting changes in the fixation target via a button-press. These changes involved a brief brightening of the target or a letter superimposed on it, each occurring probabilistically at 0.2/sec. Scores (% detected targets) were shown at the end of each run. Children were carefully monitored via an in-bore face camera and an intercom to ensure that they kept fixating and were lying still comfortably during each run, and that they were happy to continue.

### Analysis

Freesurfer (http://surfer.nmr.mgh.harvard) was used to reconstruct the cortical surface from the structural scan, and SPM8 (http://www.fil.ion.ucl.ac.uk/spm) was used to pre-process functional data. Preprocessing involved realignment, slice-time correction, and computing a co-registration matrix. Functional data was then projected onto the reconstructed cortical surface mesh, by sampling time courses from voxels midway between the white and grey matter surface. Linear trends were removed and time courses were normalized.

We used the SamSrf Matlab toolbox for population receptive field model fitting (http://dx.doi.org/10.6084/m9.figshare.1344765). As pRF model, we took a bivariate Gaussian Distribution with free parameters X, Y, and Sigma, corresponding to preferred retinotopic location and pRF size respectively. To predict the response of each pRF model to the stimuli, we integrated the bivariate Gaussian described by each model across a binarised stimulus image for each TR. The resulting time series was convolved with a standard HRF function (Haas et al., 2014) to account for delays in the BOLD response.

To identify which pRF model (which X, Y and Sigma) best predicted the measured time course, we then used a two-step procedure. (1) In a coarse fitting step, we applied heavy spatial smoothing (FWHM kernel = 8.3 mm on the spherical surface mesh) to reduce local minima. We then took a coarse, 3-dimensional search grid of the parameter space, and computed for each cortical surface vertex which parameter combination yielded the highest Pearson correlation between the measured and predicted time series. These highest-correlating parameters were used as starting point for the subsequent fine fitting step, unless R^2^ < 0.05, in which case the vertex was discarded. (2) In a fine fitting step, we used multidimensional unconstrained nonlinear minimization as implemented in the fminsearch function in Matlab, to compute the pRF parameters that minimized the difference between the predicted time course and the unsmoothed time course data. Next to X, Y and Sigma, this step also fitted a parameter Beta, accounting for signal amplitude.

We also fitted the data with a more complex Difference of Gaussian pRF model. This model has two additional free parameters next to the X, Y, and Sigma of the simple Gaussian pRF. These include the size of a second wider Gaussian (Sigma2), which was subtracted from the original positive component, and a scaling variable that determines the ratio of the two subtracted Gaussians (DoG-ratio). These added components effectively modulated the width of the positive Gaussian kernel and added a negative surround to create a “Mexican hat” shape (Zuiderbaan, Harvy, & Dumoulin, 2012).

Only vertices in which the best-fitting pRF model had a good fit (R^2^>0.1) were included in the analyses. To compute polar angle maps we took the counterclockwise angle between the positive X-axis and the polar X, Y coordinate of the best-fitting pRF model (atan (Y/X)). To compute eccentricity maps, we took the Euclidean distance from the polar X, Y coordinate to the origin at fixation (sqrt(X^2^+Y^2^)). For in-depth description of these methods see (Schwarzkopf et al., 2014). We manually delineated visual regions of interest V1-V3, V4, V3A, based on horizontal and vertical meridians in the polar angle map, following standard criteria (see Figure 1a). We also delineated area MST, but did not include it in the analysis because MST was difficult to identify in many participants and model fits were poor.

**Figure 1.**
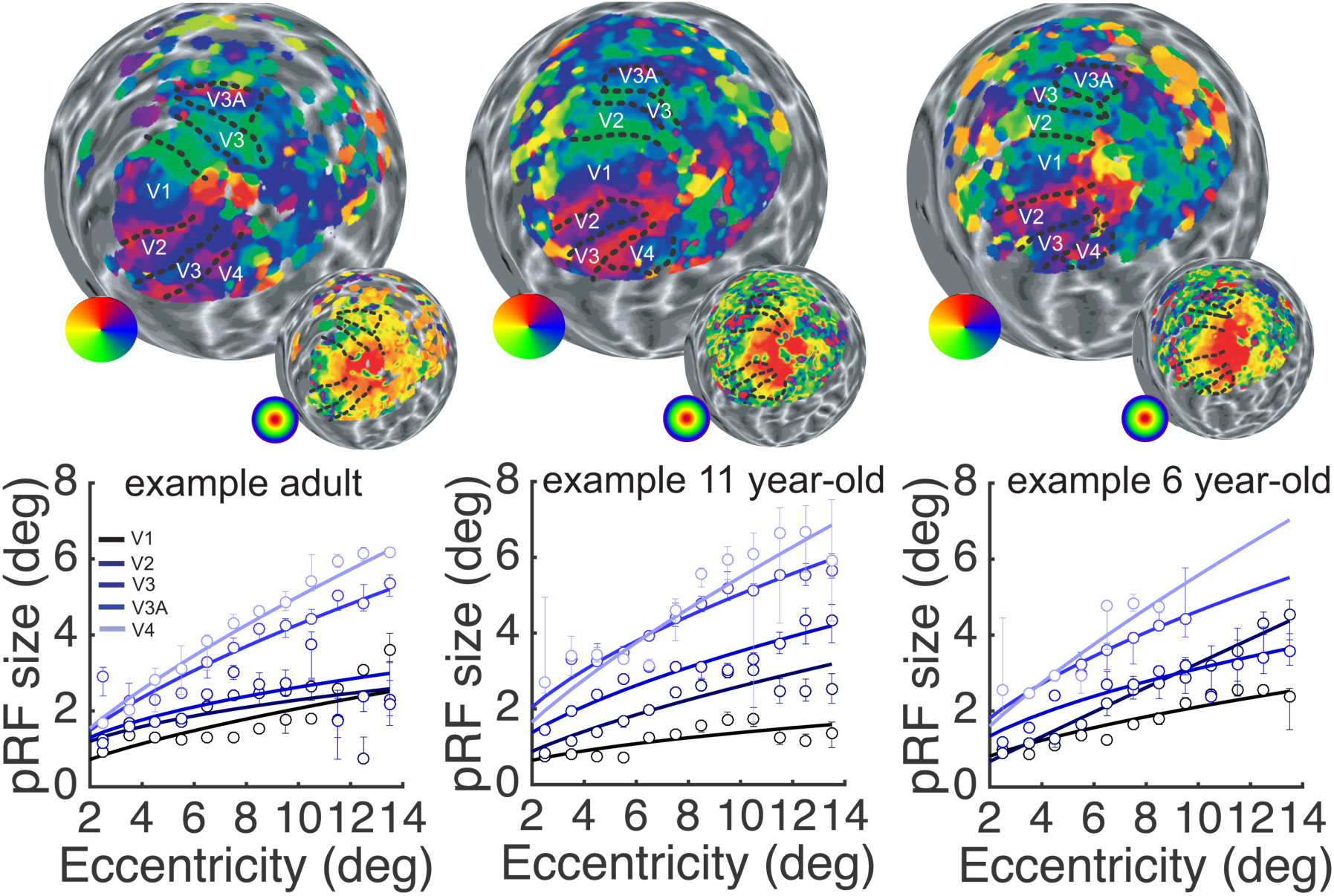
*Polar angle (large) and eccentricity (small) activation maps are displayed on the reconstructed left hemisphere (inflated to sphere) for a representative example participant in each age group. Retinotopic regions of interest defined manually based on horizontal and vertical meridians on the polar angle map are delineated. Graphs display the pRF size (the σ parameter of the best-fitting Gaussian pRF model) in degrees v/a plotted against eccentricity. Different shades of blue indicate measures from different visual areas. Circles indicated average pRF size of all vertices in the eccentricity bin with R*^*2*^*>0.1, error bars are bootstrapped 95CIs. Lines display the best fitting 2*^*nd*^ *order exponential fitted through these data points.*

## Data Quality Assurance

### Head Movement

We used stringent inclusion criteria to minimize any contributions from head-movement to any age differences in functional activation (see methods for details). After excluding participants who did not meet our criteria, there were no significant age differences in translation or rotation (translation: F(2,33)=0.14, p=0.87, rotation: F(2,33)=0.12, p=0.89, see Supplementary Figure 1).

### Fixation Task Performance

Participants were observed via an in-bore camera throughout the experiment and reminded to keep fixating when necessary. All age groups detected a high proportion of central target changes during the scanning runs (7 to 9-year-olds: 0.88 (SD= 0.05); 10 to 12-year-olds: 0.87 (SD=0.05); adults: 0.90 (SD=0.05). Thus, all participants attended well to the central marker throughout the experiment.

### Goodness of pRF model fit

Another way of ensuring that data quality was equivalent across age groups is by comparing the goodness of fit of the population receptive field model to the data. Although median Goodness of Fit (based on the Gaussian pRF model) was slightly better in adults, this difference did not reach statistical significance (F(2,34)=2.23, p=0.12). However, after removing vertices with a poorer data quality (R^2^ < 0.1) from the data, adults had slightly but significantly better model fit (F(2,34)=3.42, p=0.04).

## Results

### Population Receptive Field Size and Shape

To test for age-related changes in population receptive (pRF) size *(hypothesis i)*, we first extracted the average pRF size from each visual area of interest. PRF size was defined as the standard deviation (Sigma) of the best-fitting symmetric bivariate Gaussian. We then binned each vertex in the ROI by its eccentricity (14 bins from 1-15°) and computed the median Sigma for each bin. Only vertices with a good pRF model fit (R^2^ = 0.1) were included. Figure 1, shows delineated borders of V1-3, V4 and V3A and example data from these regions for three individual representatives of their age group. In line with previous reports, pRF size increases with eccentricity and across the visual hierarchy for each participant. This mirrors receptive field size profiles derived from single neuron recordings.

To compare how receptive field size changed across age, we took group average data (Fig 2a), and used bootstrapped ANOVA’s to test for age-differences in Sigma at each eccentricity. As the overlap in bootstrapped confidence intervals across the three different age groups indicates, there were no significant age differences in pRF size in any of the regions of interest. Out of 70 tests (5 ROI × 14 eccentricity bins), the smallest p-value was 0.059 at eccentricity bin 7-8° in V2. None of the p-values were significant at a false discovery rate of 0.05 (Benjamini and Yekutieli, 2001).

**Figure 2.**
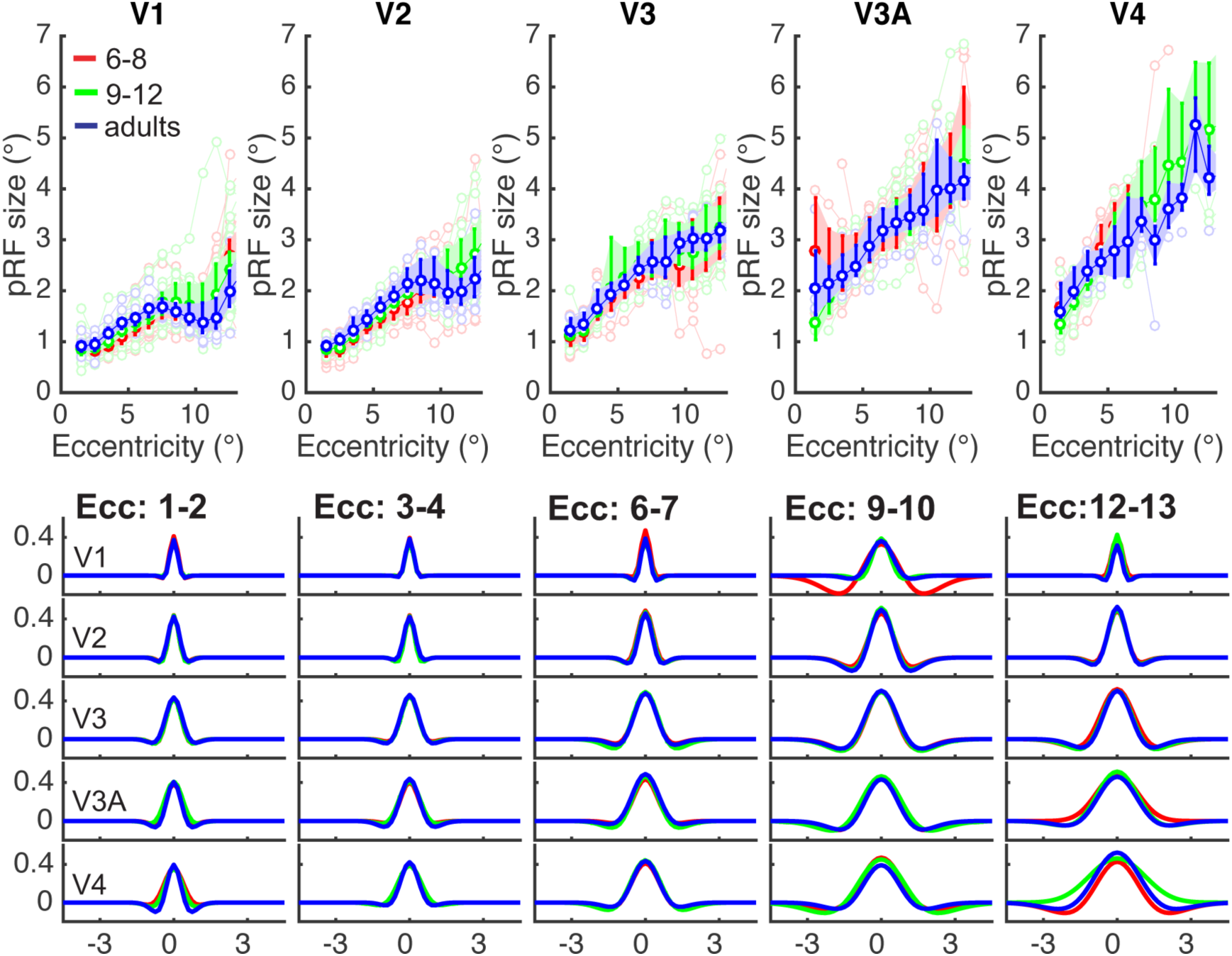
*a) average pRF size (σ) per age group plotted across eccentricity. Shaded errorbars are 95%CIs. b) Average Difference of Gaussian pRF model in each age group are plotted for 4 representative eccentricity bins, x-axis depicts width in degrees visual angle, y-axis depicts height.*

Neural receptive fields with center-surround configuration responses that are modulated by adjacent inputs are abundant in the visual system. These types of spatial dynamics are not fully captured by a Gaussian model with only an excitatory component, but are better explained by a Difference of Gaussian model with a surround inhibition component that gives it a Mexican-hat shape (Zuiderbaan, Havery, & Demoulin, 2012). Therefore, to test for age-related changes in pRF shape *(hypothesis ii)* that capture the excitation/surround suppression balance in visual cortex, we fitted this more complex pRF model to the time course data. In Figure 2b, average best-fitting DoG models are plotted across eccentricity for each age group in V1, V2, V3, V3A, or V4. As with the Gaussian pRF model, the positive kernel of the DoG pRF increases with eccentricity and along the visual hierarchy, as does the inhibitory component that models the surround modulation. Deviations from this pattern and more variability in pRF shape at larger eccentricities are likely due to interactions with receptive fields just beyond the edge of the stimulus at 14.8°, and do not indicate significant differences across age group; Bootstrapped ANOVAs comparing the three parameters of the DoG model across age per eccentricity bin revealed no significant age differences. Out of the three DoG parameters, the smallest *p*-value was 0.005. No *p*-values were significant at a false discovery rate of 0.05 (see Supplementary Figure 2). This suggests that there are no substantial changes in surround inhibition of pRFs in visual cortex between ages 7-12 years and adulthood.

### Population Receptive Field Distribution

To test for changes in cortical magnification (*hypothesis iii*), we computed the cortical magnification factor (CMF): the distance along the cortical surface required to represent a 1° distance across the visual field (Figure 3a). Analogous to the pRF size and shape analyses, we then binned each vertex with R^2^ > 0.1 by its eccentricity between 1-15°, and computed the median CMF. Bootstrapped ANOVAs comparing age groups separately for each eccentricity revealed no significant age differences in cortical magnification; The smallest out of 70 tests (5 ROI x14 bins) was p=0.012, with no p-values significant at a false discovery rate of 0.05.

**Figure 3.**
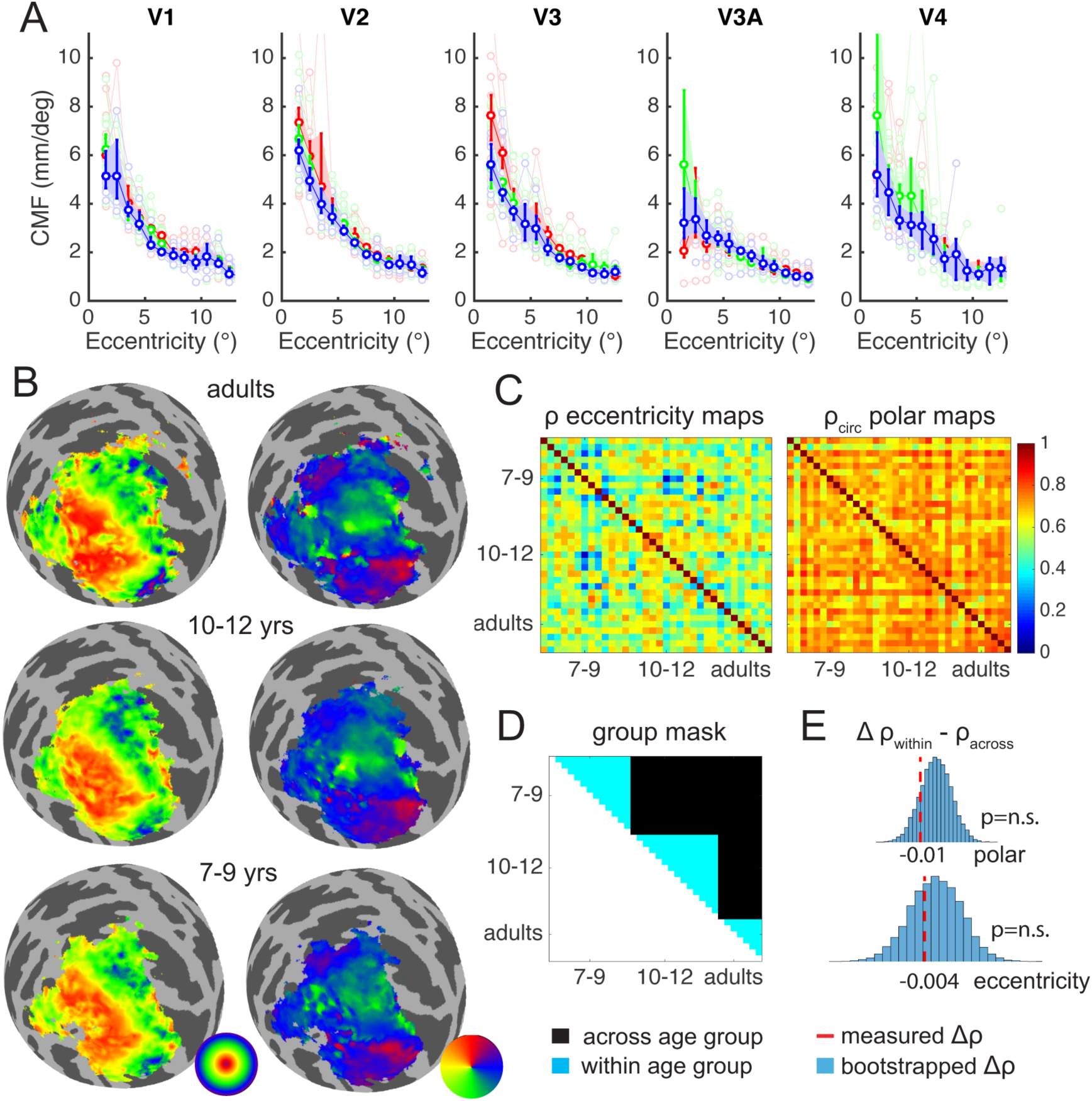
*a) Cortical magnification factor across age and visual ROI. Errorbars are bootstrapped 95%CI b) Retinotopic maps averaged across participants in each group. They are projected onto the cortical surface of the occipital lobe inflated to a sphere (left hemisphere only). Eccentricity maps (left column) and polar angle maps (right column) are shown for each age group. Only vertices with R*^*2*^ *>0.05 are included in the average. c) Matrices visualizing similarity quantified by Pearson’s correlations between eccentricity maps (left) and circular correlations between polar angle maps (right) within occipital cortex for all pairs of participants sorted by age. d) Group mask for correlation comparison. Black areas: correlations averaged to compute similarity across age groups, blue areas: correlations averaged to compute similarity within age groups. e) Red dashed lines indicate the difference in mean correlation within vs. across age groups for polar (top) and eccentricity (bottom). Negative values indicate larger correlations across groups than within. Blue H0 distributions were obtained by shuffling correlations randomly across individuals without respecting age group boundaries, and computing the mean across cells that originally contained across and within age group correlations (black and blue areas in d).*

To investigate if the overall layout of retinotopic maps was changing with age, we tested for changes in spatial consistency across individuals (*hypothesis iv*). In order to compare activation maps directly across different participants, we first aligned all individual surfaces to the Freesurfer fsaverage template so they were in a common space. We then sorted all participants by age, and calculated pair-wise Pearson’s correlations between their eccentricity maps, and pair-wise circular correlations between their polar angle maps. In Figure 3c, we have plotted the resulting correlations across the eccentricity (left) and polar angle map (right) of each possible pair of participants in two similarity matrices. Three subjects from the 10 to 12-year-old age group were excluded from this analysis, because their cross-participant map correlation (the column or row average in Fig 3c) was 3 mean absolute deviations from the median lower than for other participants. This most likely reflects poor alignment of their surface to the Freesurfer average due to movement artifacts in the structural scan. To ensure that activations included in this analysis covered the same cortex regions for or all individuals, we excluded vertices with a poor pRF model fit (excluding R^2^ < 0.05 in any of the participants). A more lenient cut-off (e.g., excluding vertices with R^2^ < 0.01 in more than 15 participants) yielded similar results.

We tested if map similarity changed with age, by taking the average correlation between retinotopic maps within age groups (blue areas in Figure 3d), and comparing it to the average correlation between maps across age groups (black areas in Figure 3d). If the spatial consistency of retinotopic maps changes across the tested age range, we should expect correlations amongst participants from the same age group to be higher than amongst those of different age groups (blue cells in Fig 3d should be “hotter” in color). This pattern is not clearly apparent when inspecting the similarity matrices in 3c, as correlations appear homogenous across all cells. Indeed, our analyses revealed a slightly higher mean correlation between individuals from *different* age groups than between individuals from the same age (Δρ_within vs. across age_ polar map = −0.01; Δρ_within vs. across age_ eccentricity map = −0.04, dotted lines fig 3e). This difference in the unexpected direction was not significant; it did not exceed the 95CI of a bootstrapped null-distribution that was obtained by shuffling correlations randomly across individuals and age groups (blue bars, fig 3e). In sum, we found no evidence for a systematic change in the distribution of population receptive tuning functions along visual cortex. Instead, cortical magnification factor and consistency of retinotopic maps were similar across children and adults.

### V1 size

Population receptive fields and cortical magnification factor are both correlated with V1 size (Harvey and Dumoulin, 2011), and visual cortex size might still increase across the tested age range (Jernigan et al., 2016). We therefore tested if any age differences in population receptive fields might be masked by systematic differences in the cortical surface area of V1 across age groups. Figure 1 shows anatomically- (left) and functionally- (right) defined V1 sizes per age group. Anatomical V1 borders were defined based on anatomical landmarks using Hinds’ probabilistic atlas (Hinds et al., 2008), as implemented in Freesurfer. Areas that belonged to V1 with an 80% probability were included in the structural label. Functional V1 borders were delineated manually based on the individual polar and eccentricity maps ((Tootell et al., 1998); see Figure 2). There were no significant age-differences in V1 size in either measure, so size differences are unlikely to have masked substantial development of population receptive fields. This is in line with findings from large-scale population studies showing that changes in occipital cortex across the tested age range are very small, in the order of ~3% (Jernigan et al., 2016). Functionally defined V1 covered a smaller area of cortex than the structurally defined V1 due to limited visual field stimulation. In line with previous findings (Dougherty et al., 2003), there was substantial (~2-fold) variability in V1 size across individuals of all ages (Fig 1, scatterplot inset; Pearson’s correlation = 0.7, *p* <0.001.

**Figure 4.**
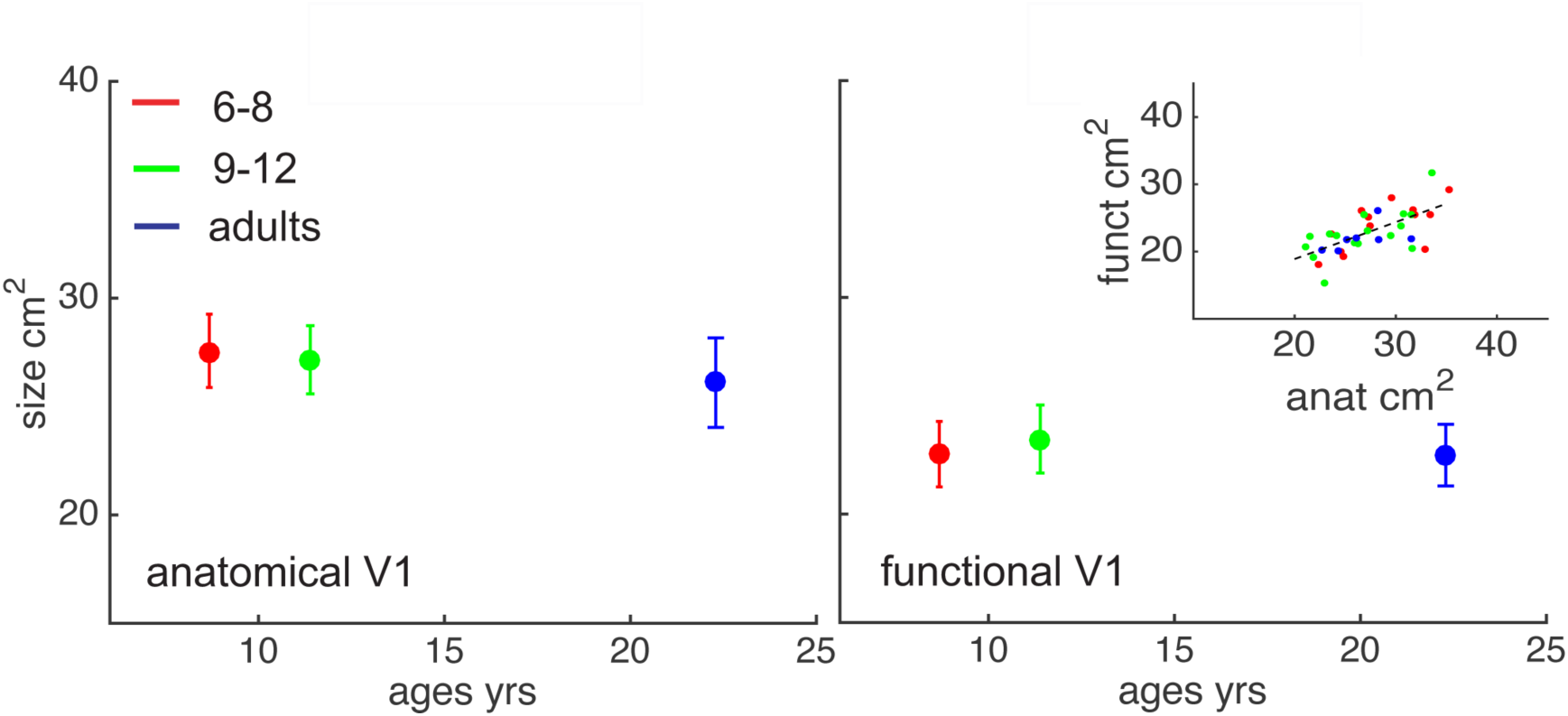
*Age differences in anatomical V1 size (left) delineated using an automated algorithm (Hinds et al., 2008), and functional V1 size (right) delineated manually using retinotopic activation maps. Errorbars are 95%CI. Right inset: correlation between anatomical and functional area size. Dotted line: best-fitting linear trend.*

## Discussion

Many basic aspects of spatial vision, including contrast sensitivity, Vernier acuity, crowded acuity, and long-range spatial integration, continue to improve until late in childhood. We used population receptive field mapping to investigate whether this can be attributed to spatial tuning of neuronal populations across occipital cortex. To do this, we delineated visual areas V1, V2, V3, V4, and V3A in groups of 6 to 9-year-olds, 10 to 12-year-olds, and adults, and compared population receptive field tuning curves and distributions within those areas. We used stringent head-movement exclusion criteria to ensure age-differences could not be explained by this confound.

To test for age differences in pRF size and shape, we fitted BOLD measures with a simple Gaussian model that varied freely in size and location, and with a difference of Gaussian model with a positive kernel and surround-suppression. For both models, pRF size increased with eccentricity and along the visual hierarchy in all participants, mirroring previous findings with adults (Dumoulin and Wandell, 2008). Moreover, we found no age differences in pRF size and surround suppression in the tested eccentricity range (up to ~15°). We also investigated differences in the distribution of pRFs along the visual field; Cortical magnification decreased from central to peripheral eccentricities in all participants, with no systematic age-differences anywhere along the visual field. Pairwise correlations of polar and eccentricity maps revealed that retinotopic organization was similar regardless of age group. In sum, our data revealed no substantial developmental changes in pRF size, shape, cortical magnification, and spatial consistency from the ages of 6-9 years onwards, suggesting spatial tuning properties of neural populations in visual cortex are close to adult-like in late childhood.

In contrast, using comparable methods, systematic differences in pRF size, cortical magnification, and surround suppression have been found in other groups with impaired visuospatial skills, including amblyopia, autism, schizophrenia, and old age (Brewer and Barton, 2014; Schwarzkopf et al., 2014; Clavagnier et al., 2015; Anderson et al., 2017). In healthy adults, the highest vs. lowest acuity at 4.7° eccentricity was linked to a 3-fold decrease in pRF size, (0.8°-2.1°; Song et al., 2015). Comparing visual performance measures and pRF properties directly across studies is difficult due to differences in methods and testing circumstances. Nevertheless, since age-related improvements in spatial vision between the age of 6 years and adulthood are often substantial with respect to variation within these age groups (*e.g.,* Carkeet et al., 1997; Skoczenski and Norcia, 2002; Jeon et al., 2010), the current finding of consistency in pRF properties across these ages is striking. It strongly suggests that the processes that limit the development of visuospatial abilities in late childhood are different from those limiting adult perception. Furthermore, the lack of evidence for prolonged tuning of neural populations up to areas V4 and V3A raises the possibility that higher-order processes beyond visual cortex may drive visuospatial improvements in late childhood. For example, children may be learning to make optimal use of the information present in visual cortex when forming their perceptual decisions.

These findings are in line with electrophysiological recordings from infant macaques suggesting that receptive field development in V1 and V2 is not the main limiting factor on visual resolution. (Kiorpes and Movshon, 2003) reported that receptive fields in young infants were adult-like with regard to orientation tuning, spatial sensitivity, and surround suppression. Receptive fields encoding the central 5° of the visual field did decrease in size (getting ~3x smaller) between the first week of life and adulthood, but the rate and time at which this happened suggested this was mainly linked to migration of cone-cells to the fovea, whilst behavioral performance improved for much longer and at much greater rates (Jacobs and Blakemore, 1988; Kiorpes and Movshon, 2003; Kiorpes, 2015). Interestingly, a similar pattern was found in MT: single neurons in MT showed adult-like tuning for speed and direction in the first 1-16 weeks after birth, even though behavioural sensitivity to random dot motion directions develops over the first 2 years of a macaque’s life. Therefore, the developmental improvement of these visual skills may be one of perceptual read-out mechanisms rather than visual cortex encoding (Kiorpes, 2015). The current results are also consistent with findings from Conner et al., (2004) who used standard retinotopic mapping methods with 9 to 12-year-old children and adults and found similar cortical magnification across these groups.

Given the close association between retinal development and receptive field size in macaques, the results of the present study suggest that the retinal mosaic may be fully established in our youngest age group of 6-9 years. Since histological data suggests that cone cell packing density and length are at only half of the adult level by 45 months (>4 years) (Yuodelis and Hendrickson, 1986), it is possible that changes in pRF tuning properties might be present at younger ages when retinal inputs to the cortex may still be reorganizing and grating acuity is still improving (Mayer and Dobson, 1982; Atkinson and Braddick, 1983; Skoczenski and Norcia, 2002). It is also possible that pRFs are still developing across our tested age in areas beyond the cortical regions that we could measure reliably. In a recent study, (Gomez et al., 2017) report evidence for increased foveal field coverage by pRFs in the fusiform face area and word form area in inferotemporal cortex. This might contribute to improved face and word perception in childhood.

If basic spatial visual skill improvements in childhood indeed depend on more efficient read-out of spatial information, this leads to the question of what might cause these changes. While beyond the scope of the current study, we may speculate that this process could be supported by age-related increases in long-range connectivity across the brain, and pruning of feedback connections (Huttenlocher and de Courten, 1987; Burkhalter, 1993; Fair et al., 2007), which could lead to greater processing speed- and noise reduction across the system, and more efficient top-down modulation (the ability to direct resources to the most informative parts of space).

It is also conceivable that some of the improvements in behavioral measures of spatial vision are in fact driven by better coping with task demands, such as improved sustained attention, reduced response bias, or more adult-like decision-bound setting. Since these factors may lead to underestimation of true performance if unaccounted for, and are more prevalent in childhood, they have been used to explain discrepancies in developmental trajectories of visual ability (Manning, Jones, Dekker and Pellicano, submitted; Leat et al., 2009). Nevertheless, behavioral measurement confounds are unlikely to account for all improvements in visuospatial performance in late childhood, because different spatial abilities develop at different rates, even when tasks are matched (Skoczenski and Norcia, 2002; Braddick and Atkinson, 2011; Kiorpes, 2015).

In conclusion, we report that population tuning in early visual cortex remains largely consistent between ages 6-9 years, 10-12 years, and adulthood. This suggests that that the development leading to improvements in basic visuospatial abilities in childhood does not entail substantial changes in cortical acuity, and is more likely due to higher-level processing such as improved neural communication, attention, or decision-making, that facilitate more optimal use of spatial information present in the visual system. Our findings highlight the importance of disentangling whether improvements in perception in childhood truly reflect improved neuronal sensitivity, or more efficient use of neuronal information. This is crucial for understanding which visual abilities develop when, which mechanisms drive this development, and the scope for plasticity at different stages of development.

## Acknowledgements

This work was supported by an ESRC Future Leaders Early Career Grant (ES/N000838/1), a Moorfields Eye Charity Grant (ST 16 03 C) awarded to TMD, and by the NIHR Biomedical Research Centre at Moorfields Eye Hospital and the UCL Institute of Ophthalmology. DSS was supported by an ERC Starting Grant (310829). BdH was supported by a DFG research fellowship (HA 7574/1-1 and /3-1).

---

To minimize age-related confounds due to head-movements in the scanner, we removed participants who made excessive sharp head-movements that are difficult to correct for using realignment (Diedrichsen and Shadmehr, 2005) from the dataset. To quantify head-movements, we computed the *absolute* rotation (mean roll, pitch and yaw in radians) and *absolute* translation (translations along the x-, y-, z- hypotenuse in mm) from each scan to the next. Participants who exceeded 1mm translation or 3° rotation from one scan to the next in more than 3 volumes in the 4 runs were excluded. To quantify head movement in the remaining participants, we summed all scan-to-scan displacements across the entire run. In Supplementary Figure 2, mean rotations and translations are plotted per individual (lighter data points) and per age group. Error bars are bootstrapped 95% confidence intervals. ANOVA’s revealed no significant age differences in translation (F(2,36)=0.131, *p*=0.88) or rotation (F(2,36)=0.086, *p*=0.91). Thus, these head-movement exclusion criteria successfully distinguished between participants who moved a lot and who moved little, resulting in age groups well matched on this potential confound.

The difference of Gaussion pRF model described below was fit to the time courses of each surface vertex following the same procedures as described in the method.

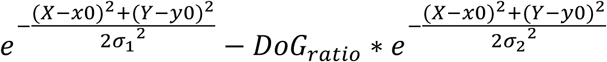

Free parameters σ1, σ2, and DoG ratio were then averaged per group and eccentricity bin. Shaded errorbars indicate the bootstrapped 95% confidence interval. Bootstrapped ANOVA’s were run separately for each eccentricity, and tested for significance at a false discovery rate of 0.05 per parameter. In line with the overlap in errorbars across the data of the three age groups in Supplementary Figure 2, this analysis revealed no significant age differences in any of the parameters: in Sigma1 (smallest uncorrected *p*=0.04), Sigma2 (smallest uncorrected *p* = 0.0058), and the DoG-ratio (smallest uncorrected *p* = 0.008).

**Supplementary Figure 1.**
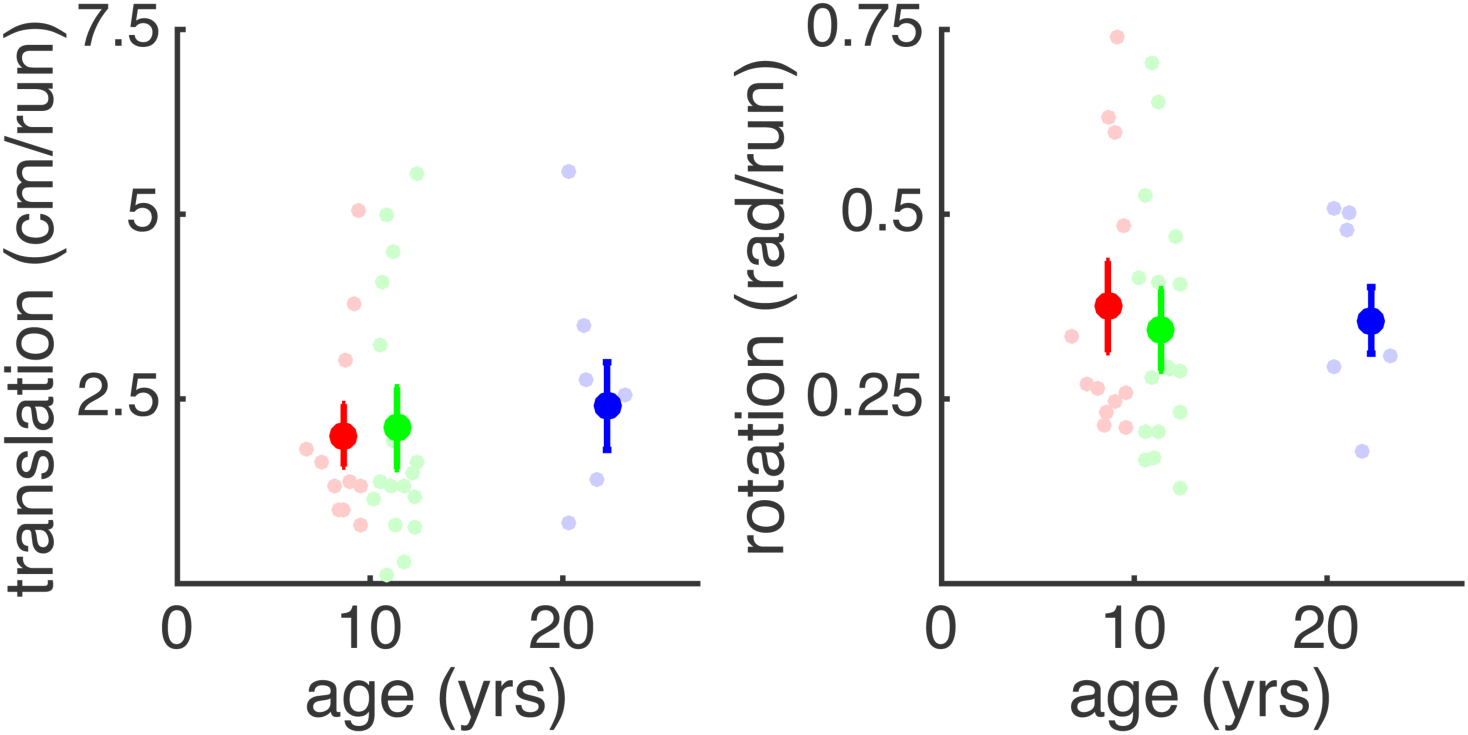

**Supplementary Figure 2.**
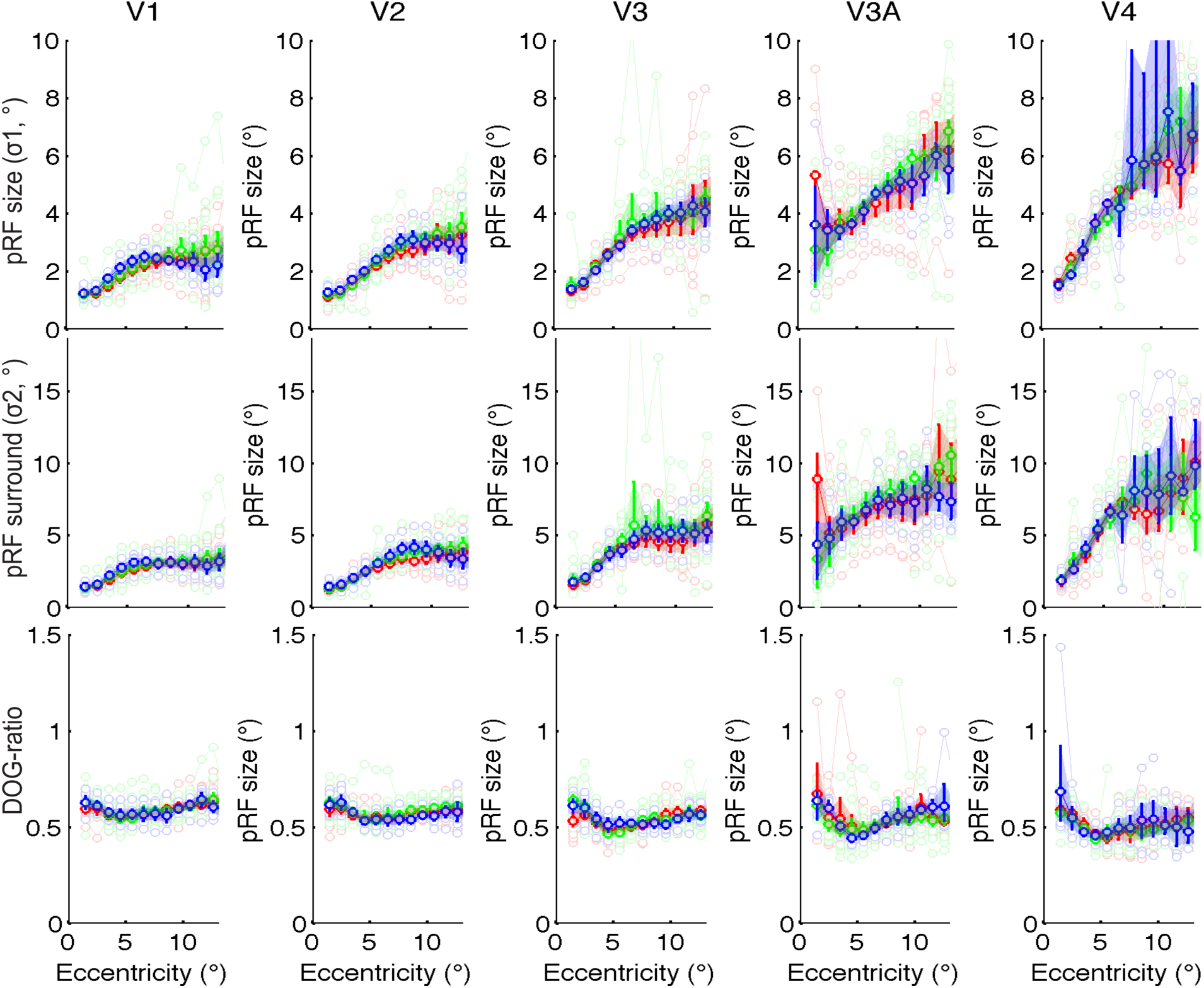

